# Low Angle Ring Illumination Stereomicroscopy (LARIS): An improved imaging method of Drosophila compound eyes

**DOI:** 10.1101/2025.07.22.666164

**Authors:** Jukta Biswas, Ankur Kumar, Anand K Singh

**Affiliations:** Department of Biology, Indian Institute of Science Education and Research Tirupati, Yerpedu, Tirupati, Andhra Pradesh, India, 517619

## Abstract

The compound eyes of Drosophila are widely used to gain valuable insights into genetics, developmental biology, cell biology, disease biology, and gene regulation. Various parameters, such as eye size, pigmentation loss, formation of necrotic patches, and disorientation, fusion or disruption of ommatidial arrays are commonly assessed to evaluate eye development and degeneration. In this study, we developed an improved optical alignment imaging technique named “Low Angle Ring Illumination Stereomicroscopy” (LARIS), which provides high-contrast images of the Drosophila compound eye. By capturing images of the same eye using different optical alignments of the stereomicroscope, we achieved the highest resolution with minimal reflection through the LARIS method. The images captured using LARIS clearly show ommatidial fusion, disorientation, and pigmentation loss in the Drosophila eye compared to those obtained with conventional imaging method. We believe that LARIS will open new avenues for improved imaging of the compound eyes of Drosophila and other insects.

---

*Drosophila* eye is comprised of approximately 750 ommatidia that are packed into a remarkably ordered three-dimensional ommatidial array covered with transparent cuticles secreted by cone cells^1^. This layer scatters light, which makes it challenging to capture high-contrast images of eyes using a stereomicroscope equipped with a standard illumination system. Analysis of such images by a trained researcher or an automated image analysis system can only give an approximation with the possibility of misinterpretation^2,3^. Scanning Electron Microscopy (SEM) is a popular technique for high-resolution imaging of the eye, but it requires imaging skills and resources^4,5^. Furthermore, SEM images provide no information on alteration in pigmentation in the adult eye. Nail polish imprint is another high-contrast imaging technique to image the molds of the fly eye, but this technique requires hand skill and does not provide pigmentation details^6,7^. A comparative analysis of different approaches used in Drosophila eye imaging is mentioned in Table 1.

**Table 1.** Comparison of LARIS with other imaging methods of adult *Drosophila* eye.

| **Features** | **Conventional** | **SEM** | **Nail Polish** | **LARIS** |
| --- | --- | --- | --- | --- |
| **Resolution** | Low | Very high | High | High |
| **Pigmentation record** | Yes | No | No | Yes |
| **Sample processing** | None | Yes | Yes | None |
| **Live/dead fly** | Live | Dead | Dead | Live |
| **Skills and resources** | Low | High | Low | Low |

## LARIS Imaging

Here, we report a simple optical alternative alignment named “Low Angle Ring Illumination Stereomicroscopy” (LARIS), that permits an inexpensive approach to obtain high-contrast imaging of the Drosophila eye using live flies under a stereomicroscope. The anesthetized and immobilized living fly is mounted sideways on a slide coated with double-sided tape so that one of the fly’s eyes is fully visible under the stereomicroscope. We tested different light sources, including a bar light, gooseneck light, diffused/straight backlight, and ring light to illuminate the eye under a stereomicroscope. Images of bar or gooseneck light illuminated eyes had considerable glare with shaded regions (Figure 1A). The dome light alignment, which provided even and diffused light, generated less glare with slightly improved image resolution (Figure 1B). The transmitted light source provided some structural details, but the resolution was poor (Figure 1C). The ring illuminator fitted around the objective lens of the stereomicroscope provides a cylinder of light illuminating the eye from top. It generates an image with no unwanted shades but an intense glare on the ommatidia (Figure 1D). However, the ring illumination images provided better resolution but lacked details of the ommatidial structure. We removed the ring illuminator from the objective lens assembly and positioned it closer to the fly on the Stereobinocular stage.

**Figure 1.**
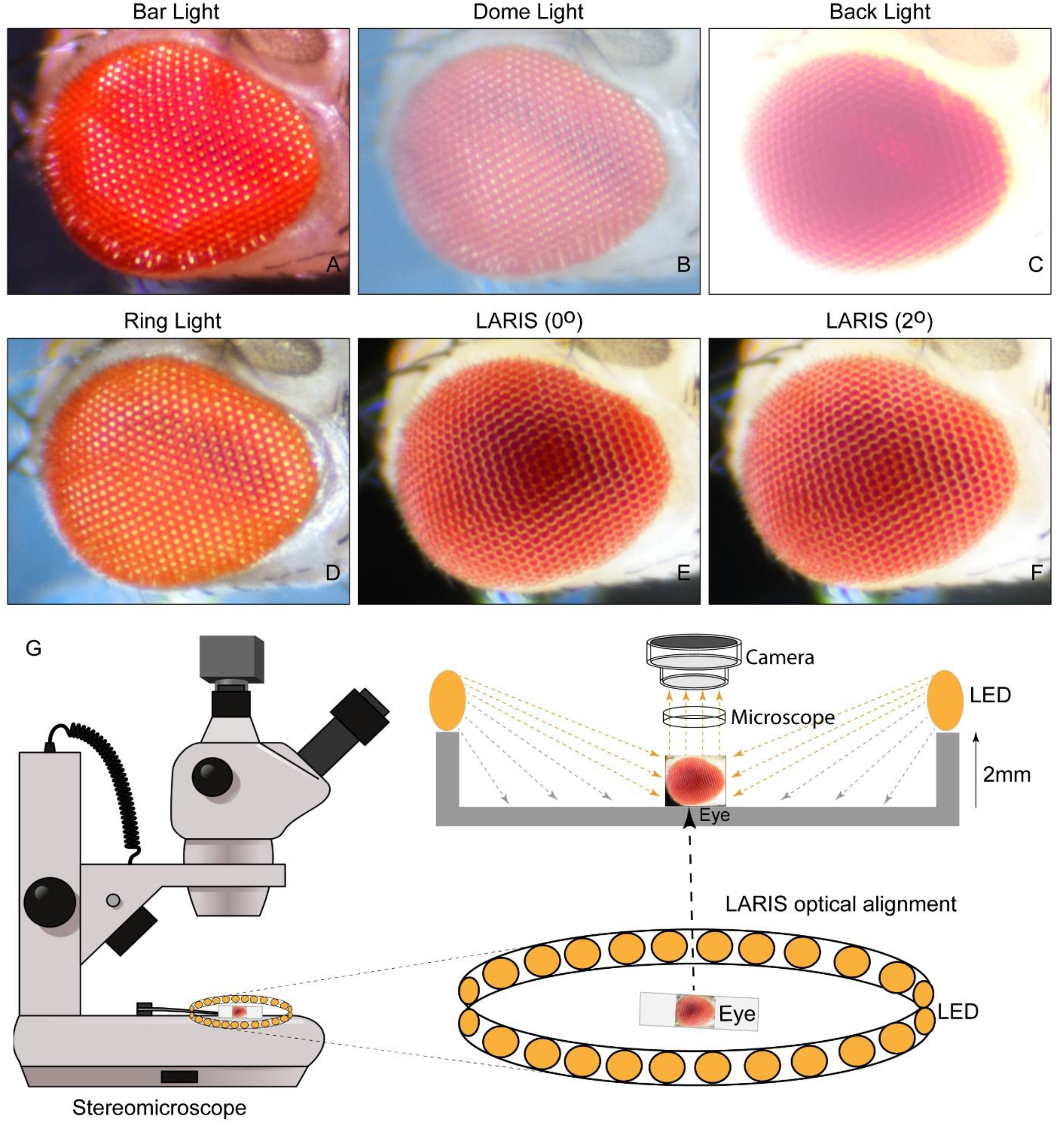
(**A-F**) Microphotographs of an adult Drosophila eye under a simple stereomicroscope with different illumination methods mentioned above each panel. (**G**) Schematic of the LARIS optical alignment for eye imaging of Drosophila. The Drosophila eye facing up is placed on a glass slide in the center of the ring illuminator. The light hits the eye from the top at ∼2^0^ to 3^0^ angle to generate a three-dimensional image of individual ommatidia without any glare. Since the illumination is at a slight angle of the uniform ring of light, a high-contrast image of the entire eye is generated. A Nikon DS-Fi3 microscope camera attached to the SMZ-745T stereomicroscope was used to capture the images at 5X magnification.

The glare disappeared when the light source and fly were at the same plane, and a high-contrast ommatidial structure appeared (Figure 1E). However, the central part of the eye has lost its contrast because of lesser light than on the peripheral regions of the curved eye surface. This issue was resolved by lifting the ring illuminator ∼2^0^ upward to the eye position. This improved optical alignment, which we call “LARIS”, did not generate any glare but illuminated all parts of the eye equally and thus provided a high-contrast image of the entire eye. (Figure 1F). The position of the ring light source is only slightly above the LARIS alignment so that depending upon the surface features of the eye, the light is scattered, and the camera receives only the scattered light from the curved surface of the eye (Figure 1G). This makes the top surface of each ommatidium slightly brighter than its basal areas, which creates a high-contrast between neighboring ommatidia and allows distinct visualization of each ommatidium (Figure 1G). The LARIS method provides better resolution of ommatidial structure, orientation and the pigmentation status compared to the conventional imaging methods of Drosophila compound eye.

We compared the conventional and LARIS imaging methods to examine the difference in structural details of normal and flies with neurodegeneration in eyes. We used *the GMR-GAL4*^8^ fly line as a control and *GMR-GAL4>PolyQ*^9^ and *GMR-GAL4>FUS*^10^ as the Huntington’s and ALS disease models (Figure 2). Under conventional imaging, the wild-type eye has only an array of bright dots without enough structural details (Figure 2A, A’). As noted above, the same eye under LARIS shows distinct ommatidia aligned in rows (Figure 2D, D’). In neurodegeneration lines where the conventional method provides the image with no distinct information on ommatidial structure (Figure 2B, B’, C, C’), LARIS images showed ommatidial fusion in *GMRGAL4>PolyQ* flies (Figure 2E, E’) and an apparent loss of ommatidial structure at the central part of the *GMRGAL4>FUS* flies (Figure 2F, F’). The abnormalities in ommatidial structure and arrangement in degenerating eyes are even more distinct in a magnified view of the LARIS image (Figure 2C, F, I).

**Figure 2.**
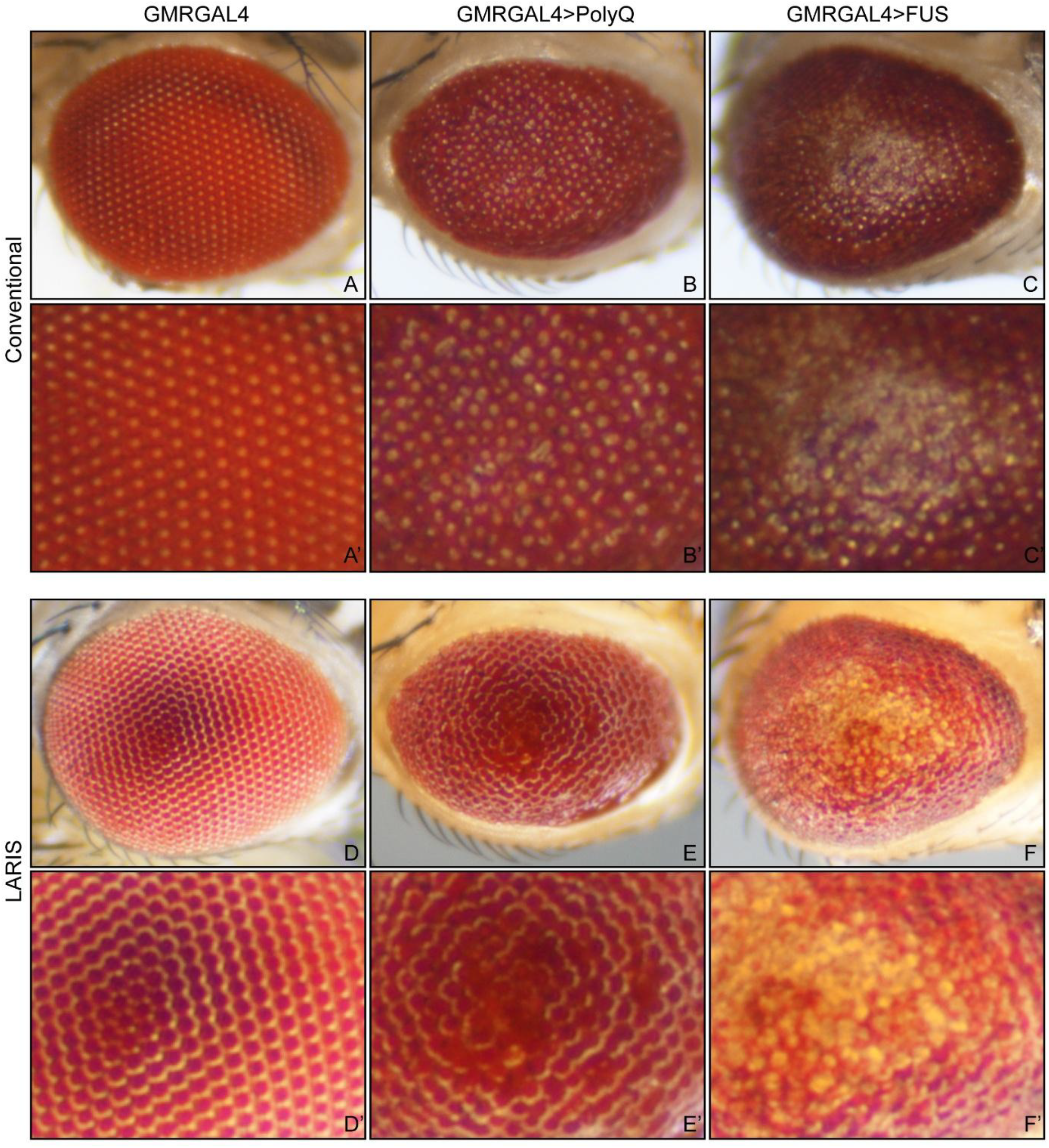
Microphotographs of adult Drosophila eye of *GMR-GAL4* (**A-C**), *GMR-GAL4> PolyQ* (**D-F**) and *GMR-GAL4>FUS* (**G-I**) using conventional (**A-C**) or LARIS imaging at the same magnification (5X) (**D-F**). A magnified view of conventional images (A’-C’) and LARIS images (D’-F’) are shown below their respective images.

## Conclusion

Our study showed that LARIS is a straightforward, reliable, and robust method for capturing high-contrast images of insect compound eyes. This technique is particularly beneficial for imaging the three-dimensional microscopic structures with highly reflective cuticles found in Drosophila and other organisms. We believe that LARIS will create new opportunities for high-contrast imaging of insect eyes.

## Acknowledgement

We acknowledge Prof. S. C. Lakhotia from Banaras Hindu University, India, for his valuable suggestions on image properties and Dr. T. Kanagasekaran from Indian Institute of Science Education and Research Tirupati for microscope optics description. We thank Science and Engineering Research Board, Grant/Award Number: CRG/2022/002767 and Department of Biotechnology, Ministry of Science and Technology, India, Grant/Award Number: BT/RHD/35/02/2006 for funding support.

## References

1. Katz, B., and Minke, B., Drosophila Photoreceptors and Signaling Mechanisms. Front. Cell.Neurosci., 2009, 3, 2.

2. Iyer, J., Wang, Q., Le, T., Pizzo, L., Gronke, S., Ambegaokar, S.S., Imai, Y., Srivastava, A., Troisí, B.L., Mardon, G., et al., Quantitative Assessment of Eye Phenotypes for Functional Genetic Studies Using Drosophila melanogaster. G3 Genes|Genomes|Genetics., 2016, 6, 5, 1427–1437.

3. Diez-Hermano, S., Valero, J., Rueda, C., Ganfornina, M.D., and Sanchez, D., An automated image analysis method to measure regularity in biological patterns: a case study in a Drosophila neurodegenerative model. Mol. Neurodegeneration., 2015, 10, 9.

4. Wolff, T., Preparation of Drosophila Eye Specimens for Scanning Electron Microscopy. Cold Spring Harb. Protoc., 2011, 1383, 5.

5. Gu, Q., Tian, Y. and Han, J., Protocol for electron microscopy of Drosophila photoreceptor cells. STAR Protocols., 2022, 3, 101496.

6. Arya, R. and Lakhotia, S.C., A simple nail polish imprint technique for examination of external morphology of Drosophila eyes. Current science., 2006, 90, 1179–1180.

7. Dwivedi, V., Tiwary, S. and Lakhotia, S.C., Suppression of induced but not developmental apoptosis in Drosophila by Ayurvedic Amalaki Rasayana and Rasa-Sindoor. J. Biosci., 2015, 40, 281–297.

8. Singh, A.K., Abdullahi, A., Soller, M., David, A. and Brogna, S., Visualisation of ribosomes in Drosophila axons using Ribo-BiFC. Biol. Open., 2020, 8, 12.

9. Raj, K. and Sarkar, S., Tissue-Specific Upregulation of Drosophila Insulin Receptor (InR) Mitigates Poly(Q)-Mediated Neurotoxicity by Restoration of Cellular Transcription Machinery. Mol. Neurobiol., 2019, 56, 1310–1329.

10. Lanson, N.A. and Pandey, U.B., FUS-related proteinopathies: Lessons from animal models. Brain Research., 2012, 1462, 44–60.

